# Cholesterol 25-hydroxylase suppresses SARS-CoV-2 replication by blocking membrane fusion

**DOI:** 10.1101/2020.06.08.141077

**Authors:** Ruochen Zang, James Brett Case, Maria Florencia Gomez Castro, Zhuoming Liu, Qiru Zeng, Haiyan Zhao, Juhee Son, Paul W. Rothlauf, Gaopeng Hou, Sayantan Bose, Xin Wang, Michael D. Vahey, Tomas Kirchhausen, Daved H. Fremont, Michael S. Diamond, Sean P.J. Whelan, Siyuan Ding

**Affiliations:** Department of Molecular Microbiology, Washington University School of Medicine, St. Louis, MO, USA; Key Laboratory of Marine Drugs, Ministry of Education, Ocean University of China, Qingdao, China; Department of Medicine, Division of Infectious Diseases, Washington University School of Medicine, St. Louis, MO, USA; Department of Pathology and Immunology, Washington University School of Medicine, St. Louis, MO, USA; Program in Molecular Cell Biology, Washington University School of Medicine, St. Louis, MO, USA; Program in Virology, Harvard Medical School, Boston, MA, USA; Autonomous Therapeutics, Inc., New York, NY, USA; Department of Biomedical Engineering, McKelvey School of Engineering, Washington University in St. Louis, St. Louis, MO, USA; Program in Cellular and Molecular Medicine, Boston Children’s Hospital and Department of Cell Biology, Harvard Medical School, Boston, MA, USA

**Author notes:** Correspondence: Siyuan Ding.

## Abstract

Cholesterol 25-hydroxylase (CH25H) is an interferon-stimulated gene (ISG) that shows broad antiviral activities against a wide range of enveloped viruses. Here, using an ISG screen against VSV-SARS-CoV and VSV-SARS-CoV-2 chimeric viruses, we identified CH25H and its enzymatic product 25-hydroxycholesterol (25HC) as potent inhibitors of virus replication. Mechanistically, internalized 25HC accumulates in the late endosomes and blocks cholesterol export, thereby restricting SARS-CoV-2 spike protein catalyzed membrane fusion. Our results highlight a unique antiviral mechanism of 25HC and provide the molecular basis for its possible therapeutic development.

## Main Text

The novel severe acute respiratory syndrome coronavirus-2 (SARS-CoV-2), the etiological agent of coronavirus disease-2019 (COVID-19)^1, 2^, has infected millions of people worldwide and caused hundreds of thousands of deaths, with a case fatality rate as high as 5%^3^. Currently, there are no FDA approved vaccines available. In most instances, treatment is limited to supportive therapies to help alleviate symptoms. Chloroquine phosphate, hydroxychloroquine sulfate, and a polymerase inhibitor remdesivir represent the only drug products that the FDA has approved for emergency use authorization^4^, and concern exists that monotherapy would rapidly result in the emergence of resistance. There is a pressing need to identify effective antivirals as countermeasures before safe and efficacious vaccines are developed and deployed. Here, we sought to harness the host innate immune responses to inhibit SARS-CoV-2 replication. Interferons (IFNs) are a group of small, secreted proteins^5, 6^ that potently suppress the replication of many viruses through the action of hundreds of IFN-stimulated genes (ISGs)^7^. IFN and ISG levels are upregulated in SARS-CoV-2 infected cells and lung tissues from COVID-19 patients^8, 9^. Compared to SARS-CoV, SARS-CoV-2 appears to be more sensitive to the antiviral activities of IFNs^10^. SARS-CoV-2 replication is inhibited by IFN treatment in both immortalized and primary cells^11, 12, 13^. While direct IFN administration often results in adverse effects in humans^14, 15^, a targeted approach of activating the antiviral effects of specific ISGs holds promise.

To identify potential anti-coronavirus (CoV) ISG effector proteins that act at the entry or egress stages of the virus replication cycle, we utilized replication-competent chimeric vesicular stomatitis virus (VSV) eGFP reporter viruses decorated with either full length SARS-CoV spike (S) protein or SARS-CoV-2 S in place of the native glycoprotein (G)^16^. We also constructed a HEK293 cell line that stably expresses plasma membrane-localized mCherry-tagged human ACE2, the SARS-CoV and SARS-CoV-2 receptor^2, 17, 18, 19^ (**Fig. S1A**). HEK293-hACE2 cells supported 100-fold more VSV-SARS-CoV-2 replication than wild-type HEK293 cells (**Fig. S1B-D**). We recently showed robust SARS-CoV-2 infection of primary human intestinal enteroids^29^. By RNA-sequencing of these intestinal enteroid cultures, we identified the ISGs most highly and commonly induced by type I IFN (IFN-β) and type III IFN (IFN-λ). We transduced HEK293-hACE2 stable cells with lentiviruses encoding 57 of these individual ISGs and tested their ability to suppress VSV-SARS-CoV and VSV-SARS-CoV-2 replication.

Ectopic expression of AXIN2, CH25H, EPSTI1, GBP5, IFIH1, IFITM2, IFITM3, and LY6E resulted in a marked reduction (< 36%) in the infectivity of both viruses, indicated by the number of GFP infected cells (**Fig. 1A, Dataset S1**). Among these genes, IFIH1 (also known as MDA5) activates IFN signaling upon ectopic expression^20^. LY6E and IFITMs recently were reported to inhibit SARS-CoV-2^21, 22^ and thus served as positive controls for our assay. We validated the top candidates in HEK293-hACE2 cells expressing CH25H, IFITM1, IFITM2, or IFITM3 respectively (**Fig. S1E**). Consistent with our screen results, the expression of IFITM2 or IFITM3 but not IFITM1 suppressed VSV-SARS-CoV-2 infection as evident by a reduction in viral mRNA and protein levels (**Fig. 1B and S1F**). CH25H was comparable to IFITM2 and blocked virus replication at 18 hours post infection (hpi) (**Fig. 1B**).

**Fig. 1.**
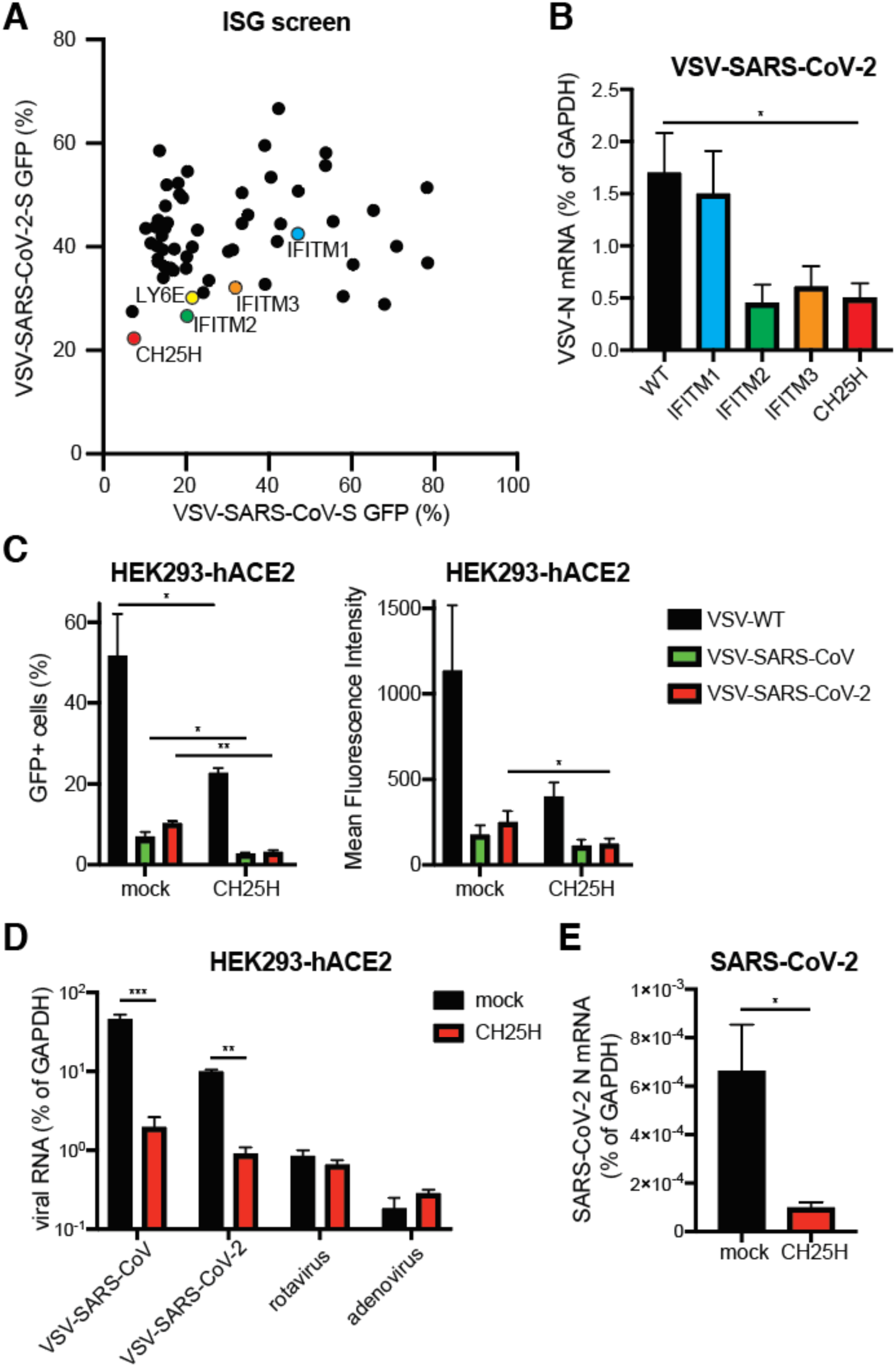
ISG screen identifies CH25H as an antiviral host factor that restricts SARS-CoV-2 infection. (A) HEK293-hACE2-mCherry cells were transduced with lentiviral vectors encoding individual ISGs for 72 hr and infected with VSV-SARS-CoV or VSV-SARS-CoV-2 (MOI=1) for 24 hr. The percentage of GFP^+^ cells were quantified and plotted. (B) Wild-type (WT) HEK293-hACE2 cells or HEK293-hACE2 cells stably expressing indicated ISGs were infected with VSV-SARS-CoV-2 (MOI=1). At 18 hpi, the mRNA level of VSV N was measured by RT-qPCR and normalized to GAPDH expression. (C) HEK293-hACE2 cells with or without CH25H expression were infected with wild-type VSV, VSV-SARS-CoV or VSV-SARS-CoV-2 (MOI=10) for 6 hr. Cells were harvested and measured for GFP percentage and intensity by flow cytometry. (D) HEK293-hACE2 cells with or without CH25H expression were infected with VSV-SARS-CoV, VSV-SARS-CoV-2, rotavirus RRV strain, or adenovirus serotype 5 (MOI=3) for 24 hr. Viral RNA levels were measured by RT-qPCR and normalized to GAPDH expression. (E) HEK293-hACE2 cells with or without CH25H expression were infected with wild-type SARS-CoV-2 (MOI=0.5). At 24 hpi, the mRNA level of SARS-CoV-2 N was measured by RT-qPCR and normalized to GAPDH expression. For all figures except A, experiments were repeated at least three times with similar results. Fig. 1A was performed twice with average numbers indicated on the graph. Raw data is listed in Dataset S1. Data are represented as mean ± SEM. Statistical significance is from pooled data of the multiple independent experiments (*p≤0.05; **p≤0.01; ***p≤0.001).

*CH25H* encodes a cholesterol 25-hydroxylase^23^ that catalyzes the formation of 25-hydroxycholesterol (25HC) from cholesterol^23^. In macrophages, 25HC is further converted to 7-α, 25-dihydroxycholesterol (7-α, 25-OHC), an oxysterol that functions as a chemoattractant for T cells and B cells^24^. 25HC exhibits broad inhibitory activities against enveloped viruses of different families^25, 26^, including two porcine CoVs^27^. Within a single-cycle of replication (6 hpi), CH25H expression inhibited the replication of VSV-SARS-CoV and VSV-SARS-CoV-2 viruses, as detected by measurement of eGFP expression using flow cytometry (**Fig. 1C**). CH25H also weakly decreased wild-type VSV replication (**Fig. 1C**), as previously reported^28^. In contrast, rotavirus and adenovirus replication were not affected (**Fig. 1D**). Unlike IFIH1, CH25H expression or 25HC treatment did not induce type I or type III IFN expression (**Fig. S1G**). The replication of a clinical isolate of SARS-CoV-2 also was suppressed in HEK293-hACE2 cells expressing CH25H compared to control plasmid transfection (**Fig. 1E**).

Next, we tested whether the antiviral activity of CH25H depends on 25HC synthesis. As compared to the control 7-α, 25-OHC, pre-treatment of HEK293-hACE2 cells with 25HC for 1 hour prior to VSV-SARS-CoV-2 infection recapitulated the suppressive effect of CH25H overexpression and reduced virus replication (**Fig. 2A**). 25HC dose-dependently inhibited VSV-SARS-CoV-2 infection in MA104 cells, with an approximate EC50 of 1.03 µM (**Fig. 2B**). No cytotoxicity was observed at the highest concentration tested (30 µM). When plaque assays were performed in the presence of 25HC, there was a reduction in both plaque numbers and sizes (**Fig. S2A-B**). Wild-type SARS-CoV-2 virus replication also was inhibited by 25HC but not 7-α, 25-OHC treatment (**Fig. 2C**). Collectively, our results suggest an antiviral activity of CH25H and its natural product 25HC in suppressing SARS-CoV-2 virus infection.

**Fig. 2.**
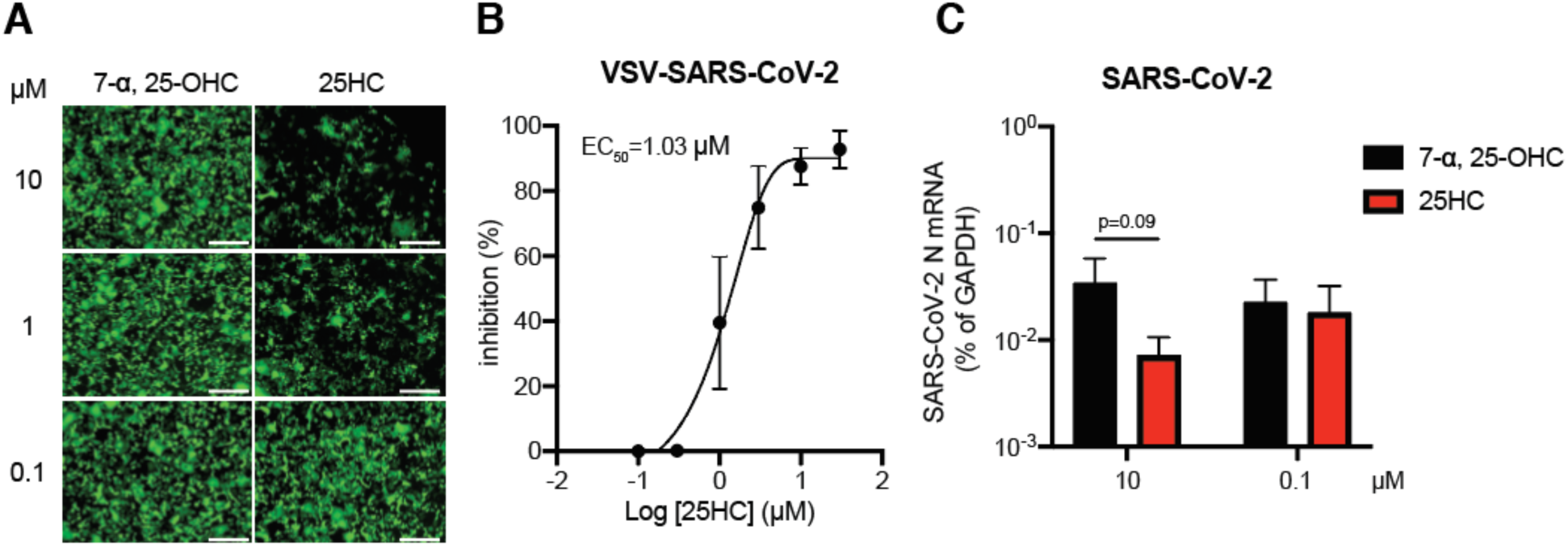
25HC inhibits SARS-CoV-2 replication. (A) HEK293-hACE2 cells were treated with 7-α, 25-OHC or 25HC at 0.1, 1, or 10 µM for 1 hr and infected with VSV-SARS-CoV-2 (MOI=5). GFP signals were detected at 24 hpi. Scale bar: 200 µm. (B) MA104 cells were treated with 25HC at indicated concentrations for 1 hr and infected with VSV-SARS-CoV-2 (MOI=0.1) for 24 hr. GFP signals were quantified by ImageJ and plotted as percentage of inhibition. (C) HEK293-hACE2 cells were treated with 7-α, 25-OHC or 25HC at 0.1 or 10 µM for 1 hr and infected with SARS-CoV-2 (MOI=0.5). At 24 hpi, the mRNA level of SARS-CoV-2 N was measured by RT-qPCR and normalized to GAPDH expression. For all figures, experiments were repeated at least three times with similar results. Data are represented as mean ± SEM. Statistical significance is from pooled data of the multiple independent experiments.

During SARS-CoV-2 entry into host cells, S protein binding to ACE2 enables its cleavage by membrane-bound TMPRSS serine proteases and subsequent fusion of the viral membrane to the host cell membrane^17, 29, 30^. Previous work suggests that trypsin treatment or TMPRSS2 expression alleviates IFITM mediated restriction of SARS-CoV and HCoV-229E entry^31, 32^. Further, TMPRSS2 is abundantly expressed in human nasal and intestinal epithelial cells^30, 33^. Thus, we examined whether the presence of TMPRSS2 assists VSV-SARS-CoV-2 to overcome ISG restriction. TMPRSS2 expression enhanced VSV-SARS-CoV and VSV-SARS-CoV-2 infection at 6 hpi (**Fig. S3A**), compared to control HEK293-hACE2 cells (Fig. 1C). Unlike IFITM3, CH25H partially retained its antiviral activity and led to reduced VSV-SARS-CoV-2 replication in TMPRSS2-expressing cells (**Fig. 3A**). Similarly, wild-type SARS-CoV-2 replication was inhibited by CH25H and 25HC in TMPRSS2 expressing cells (**Fig. 3B**).

**Fig. 3.**
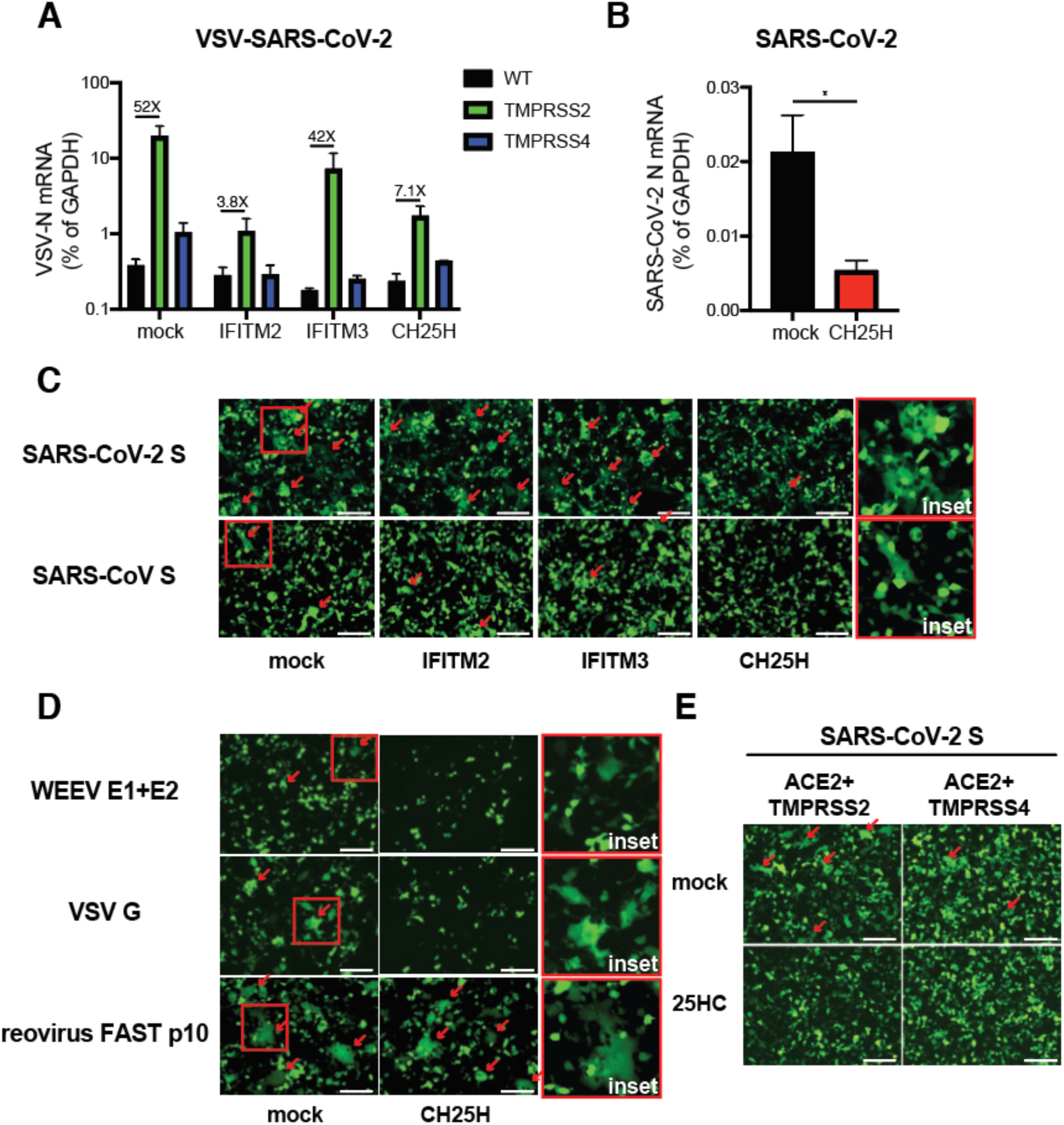
CH25H and 25HC block SARS-CoV-2 S mediated membrane fusion. (A) Wild-type (WT) HEK293-hACE2 cells or those stably expressing TMPRSS2 or TMPRSS4 were transfected with mock, IFITM2, IFITM3, or CH25H for 24 hr and infected with VSV-SARS-CoV-2 (MOI=1). At 24 hpi, the mRNA level of VSV N was measured by RT-qPCR and normalized to GAPDH expression. (B) HEK293-hACE2-TMPRSS2 cells with or without CH25H expression were infected with wild-type SARS-CoV-2 (MOI=0.5). At 24 hpi, the mRNA level of SARS-CoV-2 N was measured by RT-qPCR and normalized to GAPDH expression. (C) HEK293-hACE2-TMPRSS2 cells were co-transfected with GFP, either SARS-CoV S or SARS-CoV-2 S, and IFITM2, IFITM3, or CH25H for 24 hr. The red arrows highlight the syncytia formation. Enlarged images of mock condition are highlighted by red boxes and included as insets. Scale bar: 200 µm. (D) HEK293 cells were co-transfected with GFP, Western equine encephalomyelitis virus (WEEV) E1 and E2, VSV G, or reovirus FAST p10, with or without CH25H for 24 hr. The red arrows highlight the syncytia formation. Enlarged images of mock condition are highlighted by red boxes and included as insets. Scale bar: 200 µm. (E) HEK293-hACE2 cells stably expressing TMPRSS2 or TMPRSS4 were co-transfected with SARS-CoV-2 S and GFP with or without 25HC (10 µM) for 24 hr. The red arrows highlight the syncytia formation. Scale bar: 200 µm. For all figures, experiments were repeated at least three times with similar results. Data are represented as mean ± SEM. Statistical significance is from pooled data of the multiple independent experiments (*p≤0.05; **p≤0.01; ***p≤0.001).

We next examined the effect of 25HC on SARS-CoV S and SARS-CoV-2 S mediated membrane fusion, since 25HC blocks cell fusion by Nipah F and VSV G proteins^28^, which are class I and class III viral fusion proteins respectively^34^. We set up *an in vitro* cell-to-cell fusion assay based on the expression of S, GFP, ACE2, and TMPRSS2 in HEK293 cells, independently of virus infection (**Fig. 3C**). CH25H expression substantially reduced syncytia formation mediated by SARS-CoV-2 S (**Fig. 3C**). Although IFITM2 and IFITM3 inhibited VSV-SARS-CoV-2 replication (**Fig. 1A-B**), neither prevented S-mediated fusion (**Fig. 3C**), suggesting a distinct mode of antiviral action. Compared to SARS-CoV-2 S, SARS-CoV S induced weaker cell fusion as recently reported^35^, and this process was also blocked by CH25H expression (**Fig. 3C**). CH25H also inhibited the syncytia formation induced by Western equine encephalitis virus glycoproteins (class II) and VSV-G (class but not reovirus FAST p10 (class IV) fusion protein^36^ (**Fig. 3D**). To mimic the virus-cell membrane fusion, we co-transfected SARS-CoV-2 S and GFP into donor cells and mixed at 1:1 ratio with ACE2+TMPRSS2+TdTomato co-transfected target cells. As expected, we observed robust syncytia formation under mock conditions (**Fig. S3B**). CH25H expression in ‘recipient’ cells almost completely abolished cell-cell fusion (**Fig. S3B**). Exogenous 25HC treatment phenocopied CH25H expression and blocked SARS-CoV-2 S mediated syncytia formation (**Fig. S3B and 3E**). Similar to CH25H, 25HC failed to inhibit reovirus FAST p10 mediated fusion (**Fig. S3C**).

To define the underlying antiviral mechanisms of the IFN-CH25H-25HC axis further, we investigated whether 25HC acts on viral or host membranes. Pre-incubation of VSV-SARS-CoV-2 with 10 µM of 25HC for 20 minutes had no effect on infectivity, as opposed to the pre-treatment of host cells (**Fig. S4A**). The timing of 25HC addition suggests it primarily acted at the entry stage of VSV-SARS-CoV-2 replication (**Fig. S4B**). We examined a series of early events and excluded possible effects of 25HC on: (i) ACE2 surface levels; (ii) S cleavage by TMPRSS2; (iii) lipid raft localization, stained by a fluorophore-conjugated cholera toxin subunit B, (iv) plasma membrane fluidity, stained by 6-dodecanoyl-2-dimethylamino naphthalene (Laurdan)^37^, (v) endosomal pH, and (vi) its ability to directly bind to recombinant SARS-CoV-2 S protein (**Fig. S4C-D** and data not shown).

23-(dipyrrometheneboron difluoride)-24-norcholesterol (TopFluor-cholesterol) and [4- (dipyrrometheneboron difluoride) butanoyl]-25-hydroxycholesterol (C4 TopFluor-25HC) are chemically fluorescently labeled cholesterol and 25HC derivatives that have been used to study membrane incorporation and lipid metabolism^38^. C4 TopFluor-25HC retained its anti-VSV-SARS-CoV-2 activity (**Fig. S4E**) and blocked SARS-CoV-2 S induced syncytia formation (**Fig. 4A**), enabling us to use it as a tool to probe the antiviral mechanism of 25HC. After host cell uptake, C4 TopFluor-25HC exhibited punctate patterns and partially co-localized with lysobisphosphatidic acid (LBPA) positive late endosomes and LAMP1 positive lysosomes but not Rab4 positive early and recycling endosomes (**Fig. 4B**). Thus, we hypothesized that SARS-CoV-2 depends on endosomal trafficking to establish active replication. Consistent with this hypothesis, ectopic expression of Rab5 and Rab7 dominant negative mutants but not the wild-type proteins significantly decreased VSV-SARS-CoV-2 infection (**Fig. 4C and S4F**). However, Rab5 and Rab7 mutants did not have an additive effect with 25HC treatment (**Fig. 4C**), further suggesting that 25HC may act at these endosomal vesicles. 25HC is capable of binding Niemann-Pick C1 (NPC1) *in vitro*^39^, responsible for the egress of cholesterol from the endosomal/lysosomal compartment^40^. 25HC treatment led to an accumulation of intracellular TopFluor-cholesterol (**Fig. 4D**). 25HC failed to inhibit the replication of VSV-SARS-CoV-2 in serum-free media, in which the infectivity was markedly enhanced (**Fig. 4E**). Itraconazole (ICZ), a small-molecule inhibitor of NPC1 that elevates endosomal cholesterol levels^41^, mirrored 25HC and inhibited VSV-SARS-CoV-2 replication, more potently than the furin inhibitor decanoyl-RVKR-CMK (**Fig. 4F**). The antiviral activity of ICZ also depended on cholesterol and restricted VSV-SARS-CoV-2 in a cell-type independent manner (**Fig. 4F-G**). In contrast to chloroquine and camostat, both of which are antiviral but through different mechanisms, cholesterol-depleting agent methyl-β-cyclodextrin^42^ reduced SARS-CoV-2 S mediated cell-cell fusion (**Fig. 4H**). Either ICZ or 25HC also efficiently reduced syncytia formation (**Fig. 4H**). Finally, ICZ suppressed the replication of a recombinant SARS-CoV-2 virus that encodes a mNeon-Green reporter^43^ in Vero-E6 cells (**Fig. 4I**). Collectively, our data support a model that 25HC inhibits SARS-CoV-2 replication via enhancing endosomal cholesterol levels and blocking virus fusion.

**Fig. 4.**
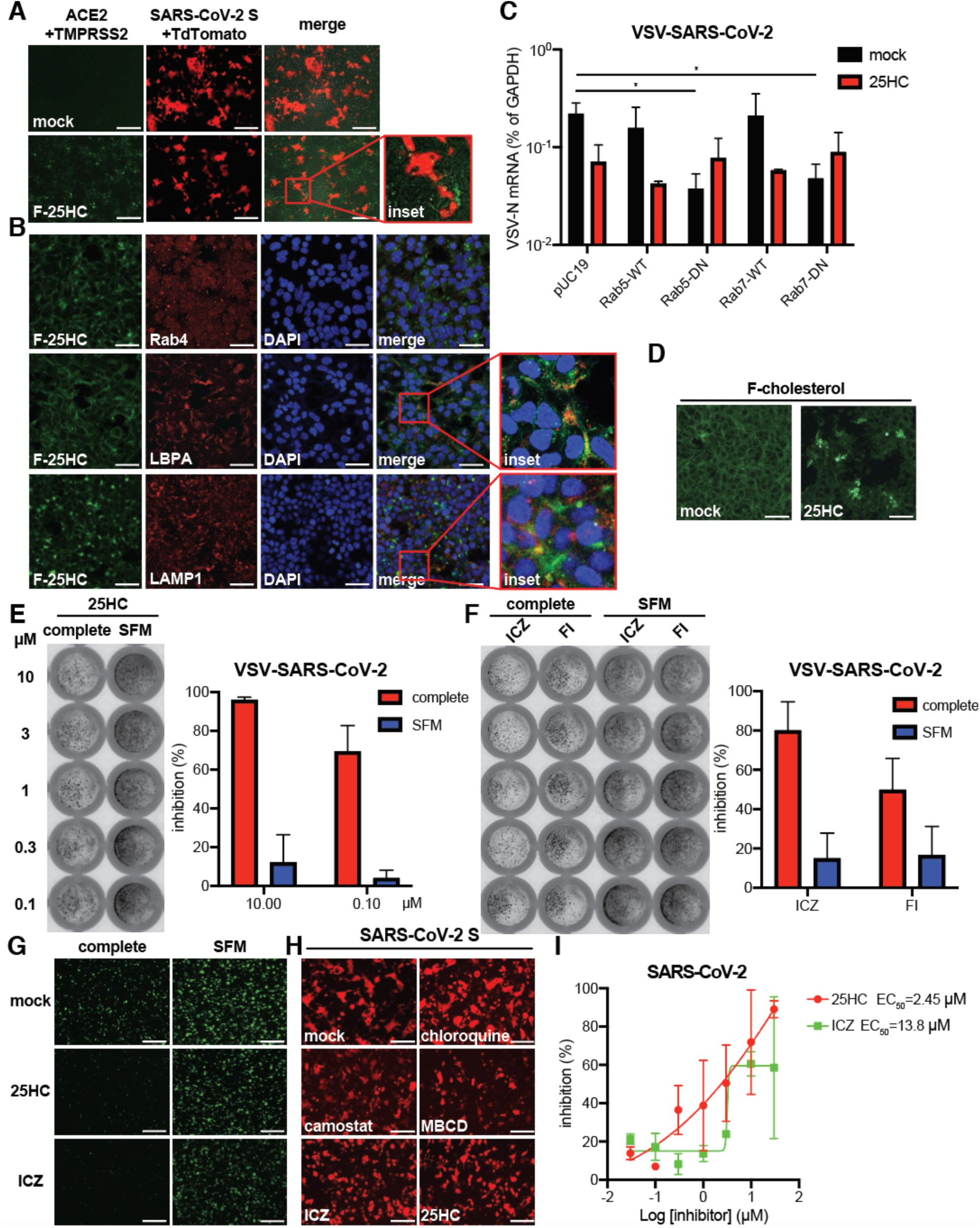
25HC inhibits endosomal cholesterol export to block SARS-CoV-2 fusion. (A) HEK293-hACE2-TMPRSS2 cells were treated with or without C4 TopFluor-25HC (F-25HC, 3 µM) and co-cultured at 1:1 ratio with HEK293 cells transfected with SARS-CoV-2 and TdTomato for 24 hr. Note that the fused cells (red) stop at the boundary of 25HC treated cells (green). Scale bar: 200 µm. (B) HEK293 cells were incubated with C4 TopFluor-25HC (F-25HC, 2 µM) for 1 hr, fixed, and stained for early/recycling endosome (Rab4), late endosome (LBPA), lysosome (LAMP1), and nucleus (blue, DAPI). Scale bar: 30 µm. (C) HEK293-hACE2 cells were transfected with wild-type (WT) or dominant negative (DN) mutants of Rab5 or Rab7 for 24 hr and infected with VSV-SARS-CoV-2 (MOI=1) with or without 25HC (10 µM). At 24 hpi, the mRNA level of VSV N was measured by RT-qPCR and normalized to GAPDH expression. (D) HEK293 cells were treated with TopFluor-cholesterol (F-cholesterol, 2 µM) with or without 25HC (20 µM) for 1 hr. Scale bar: 30 µm. (E) MA104 cells were treated with 25HC at indicated concentrations in either complete or serum-free media (SFM) for 1 hr and infected with VSV-SARS-CoV-2 (MOI=1) for 24 hr. Cells were fixed and scanned with Typhoon. Green signals were quantified by ImageJ. (F) MA104 cells were treated with itraconazole (ICZ) or furin inhibitor (FI) decanoyl-RVKR-CMK at indicated concentrations in either complete or serum-free media for 1 hr and infected with VSV-SARS-CoV-2 (MOI=1) for 24 hr. Cells were fixed and scanned with Typhoon for green signals. (G) HEK293-hACE2-TMPRSS2 cells were treated with 25HC (10 µM) or ICZ (3 µM) for 1 hr and infected with VSV-SARS-CoV-2 (MOI=1) for 20 hr. Scale bar: 500 µm. (H) HEK293-ACE2-TMPRSS2 cells were transfected with SARS-CoV-2 S and TdTomato plasmids for 24 hr in the presence of chloroquine (10 µM), camostat (10 µM), methyl-β-cyclodextrin (MCBD, 1 mM), ICZ (3 µM), or 25HC (20 µM). Scale bar: 200 µm. (I) Vero-E6 cells were treated with ICZ or 25HC at indicated concentrations for 1 hr and infected with SARS-CoV-2-mNeonGreen (MOI=0.5) for 24 hr. Cells were fixed and green signals were scanned with Typhoon and quantified by ImageJ. For all figures, experiments were repeated at least three times with similar results. Data are represented as mean ± SEM. Statistical significance is from pooled data of the multiple independent experiments (*p≤0.05; **p≤0.01; ***p≤0.001).

The identification of ISGs against different virus families have provided invaluable insights into both virus entry pathways and host innate immune system evolution^44, 45, 46, 47^. To date, few ISGs that restrict SARS-CoV replication have been identified: GILT^48^, IFITMs^32^, and recently published LY6E^21, 22^. Here, we present evidence that IFN-inducible *CH25H* and its natural product 25HC restrict S mediated membrane fusion and block SARS-CoV-2 entry into host cells. 25HC has shown broad antiviral activity against a wide range of enveloped viruses^26, 28, 49, 50^, and non-enveloped viruses such as reovirus^51^ and murine norovirus^52^. However, there seems to be two modes of inhibitory mechanisms involved. One requires a high micromolar concentration and more than 6 hours of pre-incubation time to be effective, in the case of reovirus^51^, pseudorabies virus^53^ and human papillomavirus-16^54^, suggesting an indirect metabolic/cellular pathway-mediated mechanism, whereas the other, which includes influenza A virus^26^, Lassa fever virus^55^, hepatitis C virus^56^, and SARS-CoV-2 (**Fig. 2**), functions at a low micromolar/high nanomolar range. Combined with the recent report that apilimod, a PIKfyve kinase inhibitor, effectively inhibits SARS-CoV-2 infection^57^, we confirm that this virus reaches late endosomal compartment for membrane fusion and access to the cytosol, at least in ACE2+TMPRSS2-cells. However, our data of endosomal cholesterol accumulation does not explain how the virus-cell fusion at the plasma membrane driven by SARS-CoV-2 S is effectively blocked by 25HC. A recent study demonstrates that 25HC treatment depletes ‘free’ cholesterol from the plasma membrane and prevents Listeria dissemination^58^. Treatment of cells with 25HC results in reduced cell surface but enhanced intracellular cholesterol levels (**Fig. 4D**). Therefore, it is plausible that 25HC acts at more than one subcellular compartment and that redistribution of cholesterol leads to the inhibition of membrane fusion. Our data also instruct potential drug combinations of 25HC in conjunction with those targeting the cytoplasmic steps of the SARS-CoV-2 replication cycle such as its main protease^59, 60^ or polymerase^61^. Further *in vivo* studies in animal models of SARS-CoV-2 infection and pathogenesis are required to establish the physiological impact of 25-HC-based drugs or compounds that modulate antiviral activities.

## Materials and Methods

### Plasmids, Cells, Reagents, and Viruses

#### Plasmids

Human ACE2 was cloned into pWPxld-DEST lentiviral vector with a C-terminal mCherry tag. CH25H, IFIH1, IFITM1, IFITM2, IFITM3, LY6E were cloned into pLX304 lentiviral vector with a C-terminal V5 tag. TMPRSS2, TMPRSS4 plasmids were used as previously described^30^. GFP-tagged Rab5 and Rab7 constructs were used as reported^62^. Codon-optimized SARS-CoV-2 S was a kind gift from Nevan Krogan at the University of California, San Franscico^63^. pCAGGS-SARS-CoV S was a kind gift of Paul Bates at the University of Pennsylvania^64^. pMIG-WEEV-IRES-GFP plasmid was generated by Z. Liu in the Whelan laboratory at the Washington University School of Medicine. PM-GFP and VSV-G plasmids were obtained from Addgene (#21213 and #12259, respectively). pCAGGS-FAST-p10 from pteropine orthoreovirus was generated in the Kobayashi laboratory^65^. pEGFP-N1 and pCMV-TdTomato were obtained from Clontech.

#### Cells

Human embryonic kidney cell line HEK293 (CRL-1573) were originally obtained from American Type Culture Collection (ATCC) and cultured in complete DMEM. Rhesus kidney epithelial cell lines MA104 cells were cultured in complete M199 medium. HEK293-hACE2-mCherry stable cell lines were cultured in DMEM with the addition of 5 μg/ml of blasticidin. HEK293 cells stably expressing ACE2 and TMPRSS2 were used as previously described^30^.

#### Reagents

25HC, 7-α 25-OHC, methyl-β-cyclodextrin, furin inhibitor decanoyl-RVKR-CMK, and trypsin were purchased from Sigma-Aldrich. C4 TopFluor-25-HC and TopFluor-cholesterol were purchased from Avanti Polar Lipids. 6-dodecanoyl-2-dimethylaminonaphthalene (Laurdan, D250), cholera toxin subunit B (C34777), and pHrodo™ AM Variety Pack (P35380) were purchased from Thermo Fisher. Itraconazole and camostat were purchased from Selleck Chemicals. Chloroquine (tlrl-chq) was purchased from Invivogen.

#### Viruses

Recombinant VSV-eGFP-SARS-CoV-2 was previously described^16^. VSV-eGFP-SARS-CoV was constructed in a similar manner (from S. Bose and S. Whelan, to be published separately). Adenovirus (serotype 5) and rotavirus (rhesus RRV strain) were propagated and used as previously described^66^. A clinical isolate of SARS-CoV-2 (2019-nCoV/USA-WA1/2020 strain) was obtained from the Centers for Disease Control and Prevention (gift of Natalie Thornburg). A mNeonGreen SARS-CoV-2 reporter virus was used as previously reported^43^. SARS-CoV-2 viruses were passaged in Vero CCL81 cells and titrated by focus-forming assay on Vero-E6 cells. Plaque assays were performed in MA104 cells seeded in 6-well plates using an adapted version of the rotavirus plaque assay protocol^67^. The plaque plates were scanned by Amersham Typhoon 5 (GE) and diameters were measured by ImageJ (NIH).

### RNA extraction and quantitative PCR

Total RNA was extracted from cells using RNeasy Mini kit (Qiagen) and reverse transcription was performed with High Capacity RT kit and random hexamers as previously described^68^. Quantitive PCR was performed using the AriaMX (Agilent) with a 25 µl reaction, composed of 50 ng of cDNA, 12.5 µl of Power SYBR Green master mix or Taqman master mix (Applied Biosystems), and 200 nM both forward and reverse primers. All SYBR Green primers and Taqman probes used in this study are listed in **Table S1**.

### Flow cytometry

HEK293-hACE2 or HEK293-hACE2-TMPRSS2 cells with or without CH25H expression were inoculated with wild-type VSV-GFP, VSV-SARS-CoV, or VSV-SARS-CoV-2 at an MOI = 10 (based on titers in Vero cells) for 1 hr at 37°C. At 6 hpi, cells were harvested and fixed in 4% PFA. GFP positive cells were determined by BD LSRFortessa™ X-20 cell analyzer and analyzed by FlowJo v10.6.2 (BD).

### Bright-field and immunofluorescence microscopy

For brightfield and epifluorescence, cultured cells were imaged by REVOLVE4 microscope (ECHO) with a 10X objective. For confocal microscopy, samples in 8-well chamber slides were fixed in 4% paraformaldehyde for 10 min at room temperature and stained as previously described^69^. Cells were permeabilized and stained with antibodies against DAPI (P36962, Thermo Fisher), LAMP1 (9091S, Cell Signaling), LBPA (MABT837, Sigma), and Rab4 (ab13252, Abcam). Stained cells were washed with PBS, whole mounted with Antifade Mountant, and imaged with a Zeiss LSM880 Confocal Microscope at the Molecular Microbiology imaging core facility at Washington University in St. Louis. Images were visualized by Volocity v6.3 and quantification was determined by ImageJ (NIH).

### Western blotting

Cell lysates were harvest in RIPA buffer supplemented with protease inhibitor cocktail and phosphatase inhibitor. Proteins were resolved in SDS-PAGE and analyzed by antibody as described (45) using the following antibodies and dilutions: GAPDH (631402, Biolegend), GFP (2555S, Cell Signaling), SARS-CoV-2 S1 (40590-T62, Sino Biological), SARS-CoV-2 S2 (40592-T62, Sino Biological), and V5 (13202S, Cell Signaling). Secondary antibodies were anti-rabbit (#7074, Cell Signaling) or anti-mouse (#7076, Cell Signaling) immunoglobulin G horseradish peroxidase-linked antibodies. Protein bands were visualized with Clarity ECL substrate and a Biorad Gel Doc XR system.

### Statistical Analysis

All bar graphs were displayed as means ± SEM. Statistical significance in data Fig. 1E, 3B, S1C, S1D, and S2B was calculated by Student’s t test using Prism 8.4.2 (GraphPad). Statistical significance in data Fig. 1B, 1C, 1D, 4C, S1G, and S4A was calculated by pairwise ANOVA using Prism 8. Non-linear regression (curve fit) was performed to calculate EC50 values for Fig. 2B and 4I. All data were presented as asterisks (*p≤0.05; **p≤0.01; ***p≤0.001). All experiments other than Fig. 1A were repeated at least three times. Fig. 1A was performed twice with average numbers indicated on the graph. The raw data is included in Dataset S1.

---

**Fig. S1.**
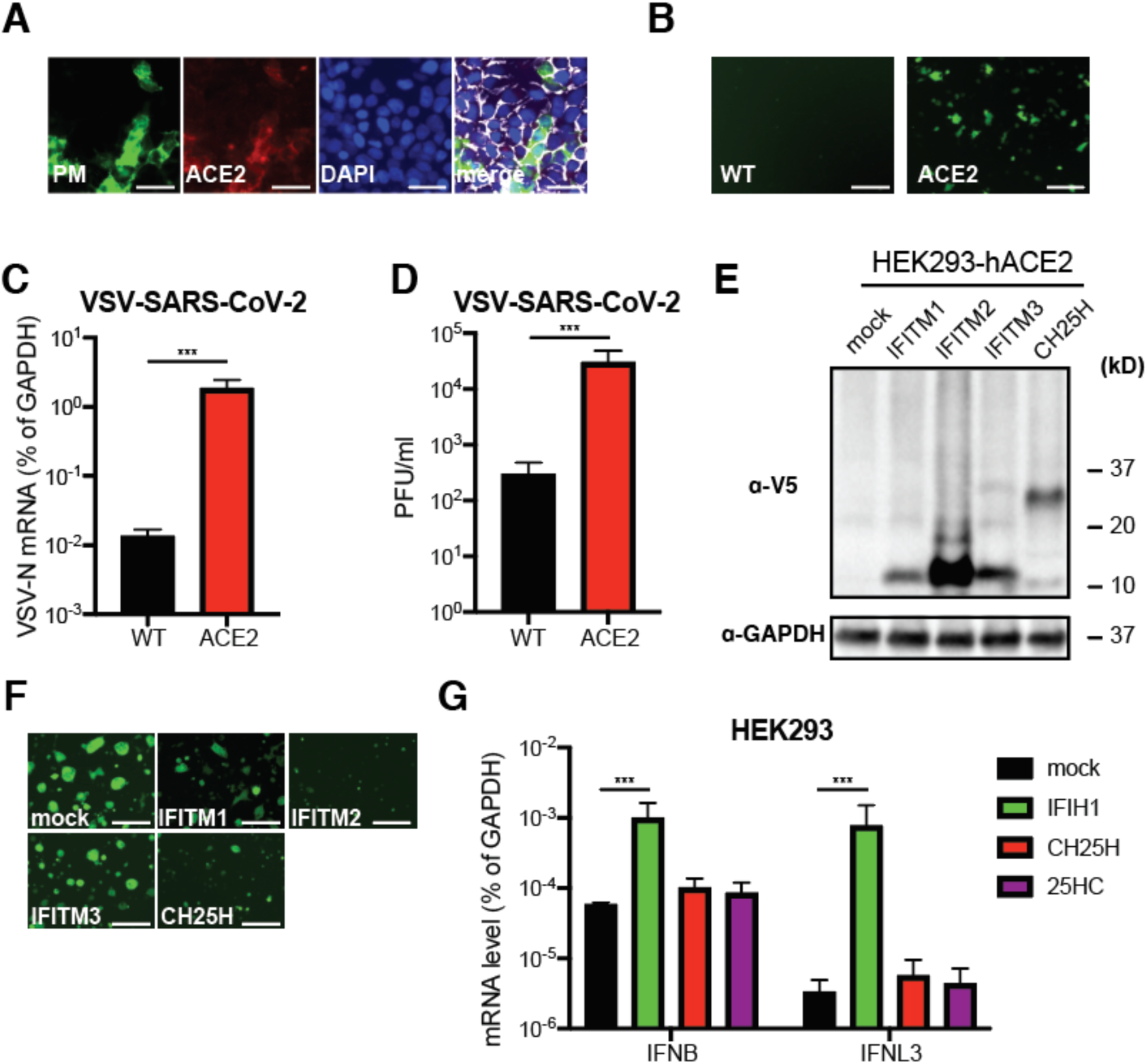
CH25H suppresses VSV-SARS-CoV-2 replication in HEK293-hACE2 cells. (A) HEK293-hACE2-mCherry cells were transfected with plasma membrane (PM)-localized GFP and stained for cell surface (green), ACE2 (red), nucleus (DAPI, blue), and actin (white). Scale bar: 30 µm. (B) Wild-type (WT) HEK293 or HEK293-hACE2-mCherry cells were infected with VSV-SARS-CoV-2 (MOI=1) for 8 hr. Scale bar: 200 µm. (C) Same as (B) except that infection was 24 hr and RNA was harvested for RT-qPCR measuring the mRNA level of VSV N compared to GAPDH expression. (D) Same as (B) except that infection was 24 hr and cell lysates were harvested for plaque assays. (E) HEK293-hACE2 cells stably expressing indicated ISGs were harvested for western blot and probed for V5-tagged ISG and GAPDH protein levels. (F) HEK293-hACE2 cells stably expressing indicated ISGs were infected with VSV-SARS-CoV-2 (MOI=1) for 24 hr. Scale bar: 200 µm. (G) HEK293 cells were transfected with mock, IFIH1, or CH25H plasmids for 24 hr or treated with 25HC (10 µM) for 1 hr. RNA was harvested and the mRNA levels of IFN-β (IFNB) and IFN-λ (IFNL3) were measured by RT-qPCR and normalized to GAPDH expression. For all figures, experiments were repeated at least three times with similar results. Data are represented as mean ± SEM. Statistical significance is from pooled data of the multiple independent experiments (*p≤0.05; **p≤0.01; ***p≤0.001).

**Fig. S2.**
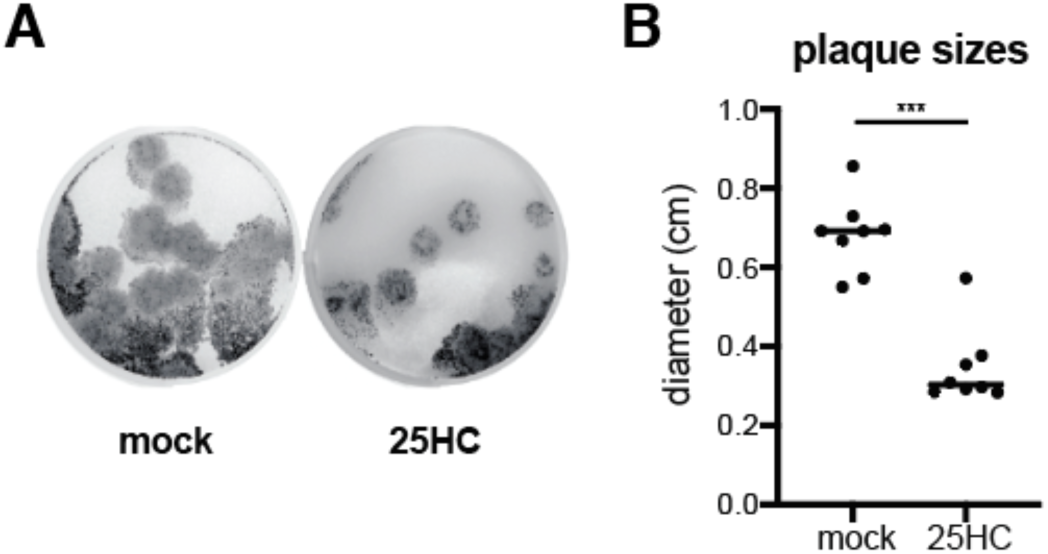
25HC restricts VSV-SARS-CoV-2 replication in MA104 cells. (A) MA104 cells were infected with serially diluted VSV-SARS-CoV-2 (10^5^ shown here) with or without 25HC (10 µM). At 3 dpi, GFP signals were scanned with Typhoon. (B) Quantification of plaque sizes in (A). For all figures, experiments were repeated at least three times with similar results. Individual data point is indicated (*p≤0.05; **p≤0.01; ***p≤0.001).

**Fig. S3.**
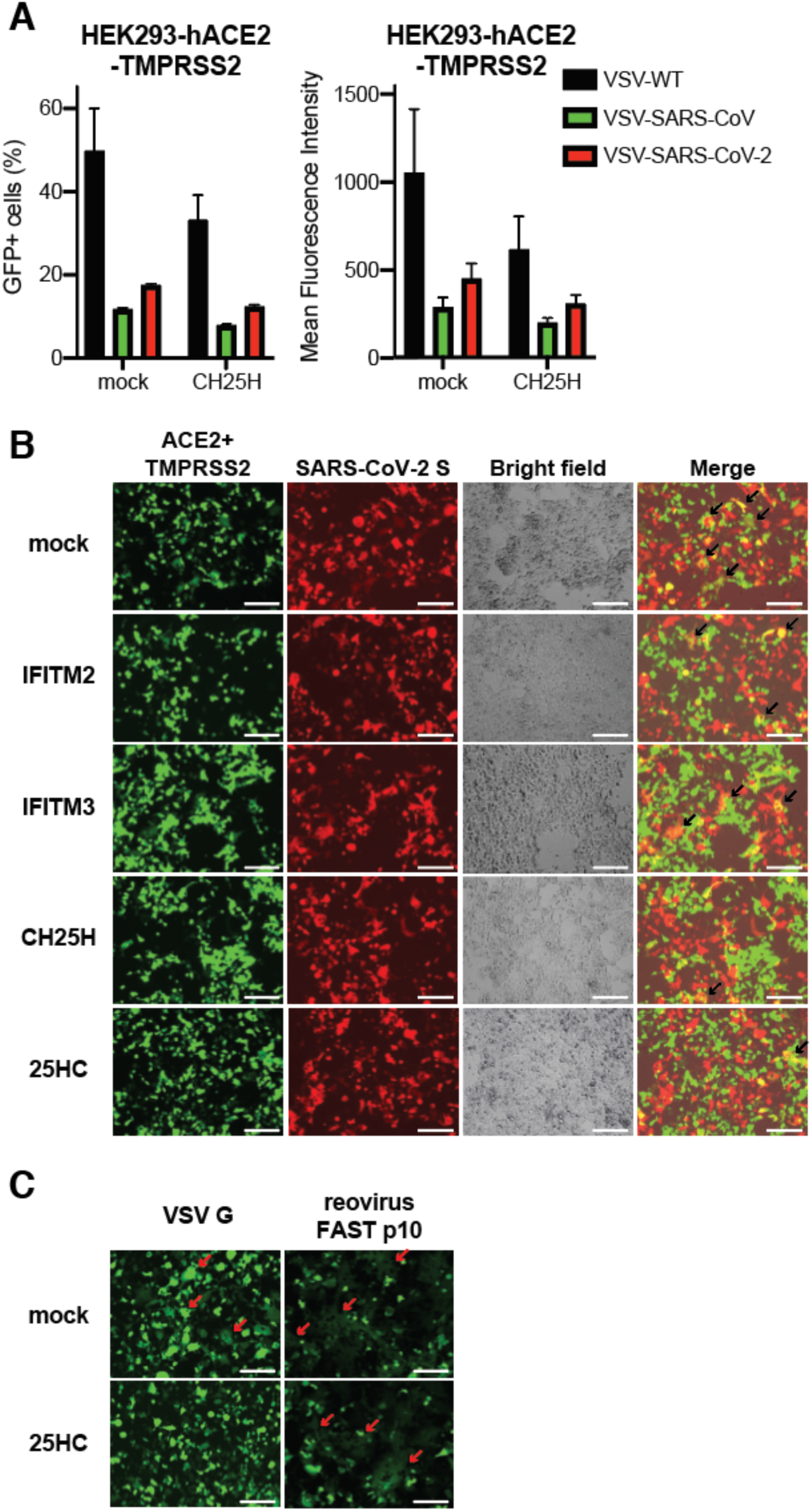
CH25H and 25HC block SARS-CoV-2 S mediated fusion. (A) HEK293-hACE2-TMPRSS2 cells were infected with wild-type VSV, VSV-SARS-CoV or VSV-SARS-CoV-2 (MOI=10) for 6 hr. Cells were harvested and measured for GFP percentage and intensity by flow cytometry. (B) HEK293-hACE2-TMPRSS2 cells expressing GFP and indicated ISGs or treated with 25HC (10 µM) were mixed at 1:1 ratio and co-cultured with HEK293 cells expressing SARS-CoV-2 S and TdTomato for 24 hr. Note the formation of cell-cell fusion (yellow), highlighted by black arrows. Scale bar: 200 µm. (C) HEK293 cells were co-transfected with GFP, VSV G, or reovirus FAST p10, with or without 25HC (10 µM) for 24 hr. The red arrows highlight the syncytia formation. Scale bar: 200 µm. For all figures, experiments were repeated at least three times with similar results. Data are represented as mean ± SEM.

**Fig. S4.**
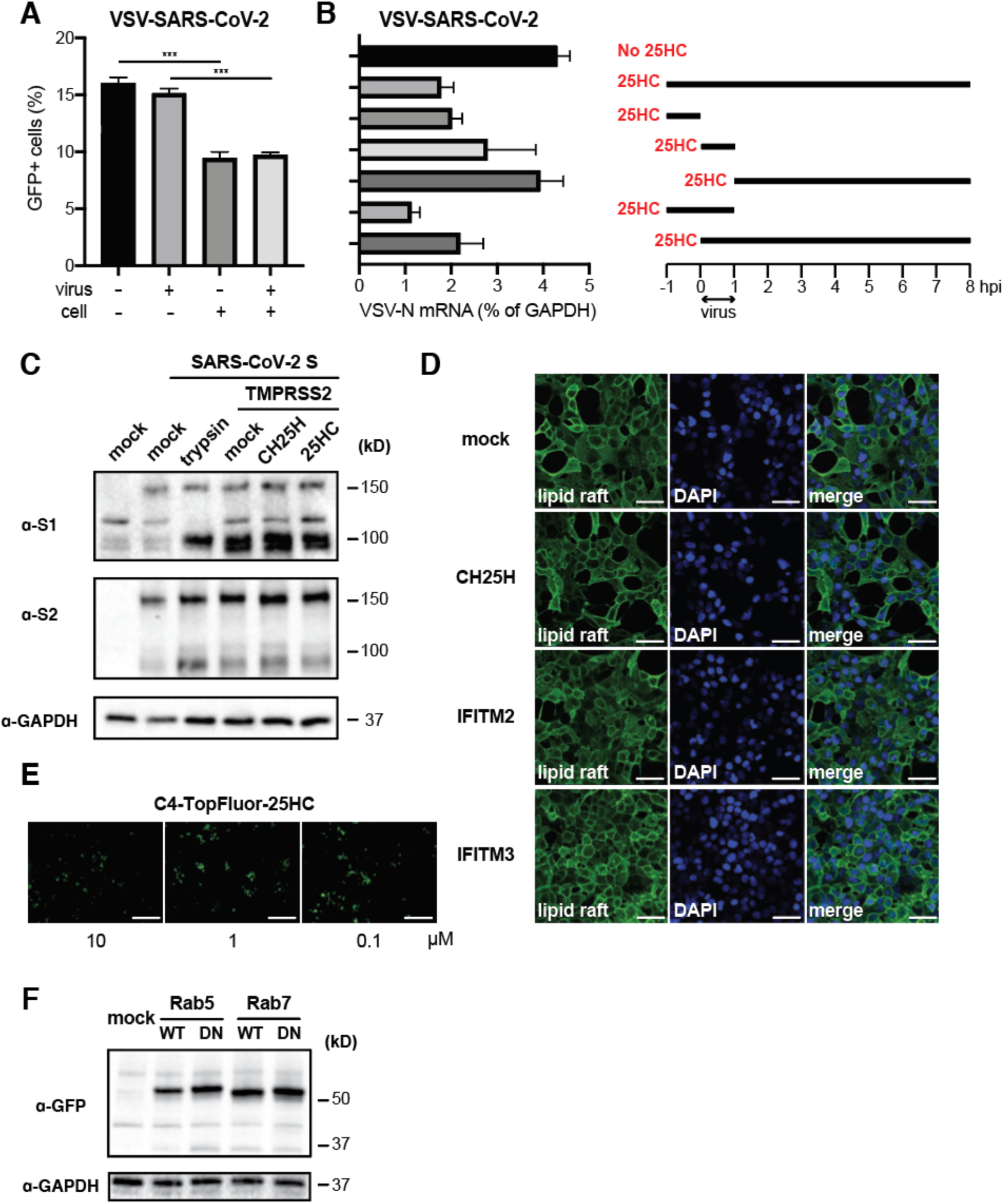
CH25H and 25HC do not affect S cleavage or lipid raft organization. (A) VSV-SARS-CoV-2 was incubated with 25HC (10 µM) for 30 min. HEK293-hACE2 cells were treated with 25HC (10 µM) for 1 hr. At 6 hpi, cells were harvested and measured for GFP percentage and intensity by flow cytometry. (B) MA104 cells were treated with 25HC (10 µM) based on the scheme (right panel) and infected with VSV-SARS-CoV-2 (MOI=1). At 24 hpi, the mRNA level of VSV N was measured by RT-qPCR and normalized to GAPDH expression (left panel). (C) HEK293-hACE2 cells were transfected with SARS-CoV-2 for 24 hr. Some cells were also transfected with TMPRSS2 or treated with trypsin (0.5 µg/ml) or 25HC (10 µM). Cells were harvested for western blot and probed for SARS-CoV-2 S1, S2, and GAPDH protein levels. (D) HEK293-hACE2 cells stably expressing indicated ISGs were stained for lipid rafts (cholera toxin B, green) and nucleus (DAPI, blue). Scale bar: 30 µm. (E) HEK293 cells were treated with C4-TopFluor-25HC (10, 1, or 0.1 µM) for 1 hr and infected with VSV-SARS-CoV-2 (MOI=0.5) for 24 hr. Scale bar: 500 µm. (F) HEK293-hACE2 cells were transfected GFP-tagged wild-type (WT) or dominant negative (DN) mutants of Rab5 or Rab7 for 24 hr. Cells were harvested for western blot and probed for GFP and GAPDH protein levels. For all figures, experiments were repeated at least three times with similar results. Data are represented as mean ± SEM. Statistical significance is from pooled data of the multiple independent experiments (*p≤0.05; **p≤0.01; ***p≤0.001).

## Acknowledgements

We appreciate the helpful discussion with Drs. Rohit Pappu (School of Engineering), Kartik Mani, Abhinav Diwan (Center for Cardiovascular research), Anil Cashikar, Steven Paul (Department of Psychiatry), and David Holtzman (Department of Neurology). We are thankful to assistance from Matthew Williams (Molecular Microbiology Media and Glassware Facility), Wandy Betty (Molecular Microbiology Imaging Facility), and Marina Cella and Erica Lantelme (flow cytometry core facility, Department of Pathology and immunology). SARS-CoV-2 Taqman probe and viral RNA standards were prepared by Adam Bailey (Division of Infectious Diseases).

## Supplemental Information

Table S1. Quantitative PCR primer information

Dataset S1. Results of ISG screens against VSV-SARS-CoV and VSV-SARS-CoV-2

